## Supplementary document for "Light-evoked deformations in rod photoreceptors, pigment epithelium and subretinal space revealed by prolonged and multilayered optoretinography"

#### This PDF file includes:

Supporting text
Figs. S1 to S3
Tables S1 to S3
Legend for Movie S1
SI References

#### Other supporting materials for this manuscript include the following:

Movie S1

### Supporting Information Text

**Estimation of the hydraulic conductivity and water permeability coefficient of the rod OS membrane.** We estimated the hydraulic conductivity and water permeability coefficient of the rod OS membrane based on a model developed by Zhang *et al.* (1). Some parameters used in this model are summarized in Table S1. In brief, the model is based on osmotic swelling of the rod OS, which occurs in response to the incremental osmotic pressure triggered by the phototransduction cascade. Assuming that the saturated elongation is purely an osmotic equilibrium, the Van't Hoff's law indicates that the volume increase is proportional to the incremental osmotic pressure,

$$\Delta V_{\text{cyto}}/V_{\text{cyto, rest}} = \Delta\Pi/\Pi_{\text{rest}}, \tag{S1}$$

where $\Pi_{\text{rest}}$ and $V_{\text{cyto, rest}}$ are the osmotic pressure and the cytoplasmic volume of the rod OS in the resting (dark-adapted) state. $\Delta\Pi$ is the phototransduction-induced increase in osmotic pressure, $\Delta V_{\text{cyto}}$ is the incremental cytoplasmic volume when a new osmotic equilibrium is established. Since a moderate osmotic perturbation does not alter the rod OS width and the disc membranes occupy ∼50% of the rod OS volume, the fractional volume increase $\Delta V_{\text{cyto}}/V_{\text{cyto, rest}}$ can be estimated by $2\Delta L/L$ (1). As a result, our measurements in Fig. 4B indicated that a flash with a bleach level of 0.28% increased the rod OS cytoplasmic osmolarity $\Delta\Pi/RT = 0.91$ mOsM. Note that this value should be considered as a lower limit, because the restoring elastic force from expansion is ignored.

Considering the start of the phototransduction cascade, the water inflow rate is determined by the initial increase of osmotic pressure and the hydraulic conductivity $L_p$ of rod OS membrane,

$$J_w = L_p S_{\text{ROS}} \Delta\Pi, \tag{S2}$$

The initial water influx rate $J_w$ can be calculated from the slope of the rod OS elongation. From our measurements, a 1-ms flash with a bleach level of 0.28% yielded an initial water influx rate of $8.5 \times 10^{-2}$ µm·s$^{-1}$ in length and 0.13 µm$^3$·s$^{-1}$ in volume ($J_w$) with the cross-sectional area of 1.54 µm$^2$. Our measurements yielded a hydraulic conductivity of 4.2 µm$^3$·s$^{-1}$ and accordingly, a water permeability coefficient of $5.9 \times 10^{-3}$ cm·s$^{-1}$, which agrees with the in-vitro experiment results of $2.6 \times 10^{-3}$ cm·s$^{-1}$ measured by Preston *et al.* (2) and $2.0 \times 10^{-3}$ cm·s$^{-1}$ measured by Korenbrot *et al.* (3). It should be noted that because the effect of restoring elastic force was ignored in the above model, the estimated hydraulic conductivity and water permeability coefficient should be considered the upper limits.

**Visible-light, spectral-domain optical coherence tomography.** A visible-light optical coherence tomography (vis-OCT) was constructed using a supercontinuum laser covering 400-2300 nm spectrum (SuperK Extreme, NKT Photonics, Denmark) as the light source. A spectrum ranging from 435 nm to 650 nm with a full-width at half-maximum bandwidth of 120 nm was selected by a spectral splitter (SuperK SPLIT, NKT Photonics, Denmark) and a shortpass filter (#47-290, Edmund Optics, USA), and further reshaped by a bandpass filter (#16-362, Edmund Optics, USA). In the sample arm, a customized achromatizing triplet lens was placed between the reflective collimator and the galvo scanner (Saturn5, ScannerMax, USA) to reduce the chromatic aberration of the rat eye. A lens-based afocal telescope conjugated the center of the galvo scanner pair to the pupil with a magnification of 0.18 (scan lens: 75 mm focal length; ocular lens: 30 mm + 25 mm focal length) to reduce the incident beam diameter to 310 µm at the pupil.

**Calculation of rhodopsin bleaching.** Given the negligible regeneration of rhodopsin during a short light stimulus, rhodopsin bleaching can be calculated as following (4, 5):

$$p(Q) = \exp\left(-Q/Q_e\right), \tag{S3}$$

where $p$ is the fraction of unbleached rhodopsin present after the light stimulus, $Q$ is the energy density of the 500 nm stimulus (photons/µm$^2$), $Q_e$ is the energy density that reduces the fraction rhodopsin present to 1/e of its dark-adapted level, which was measured in rats to be $7.94 \times 10^7$ photons/µm$^2$ (4). The energy density $Q$ can be calculated as,

$$Q = \frac{Pt}{AE_v}, \tag{S4}$$

where $P$ and $t$ are the power and duration of the light stimulus, $A$ is the illuminated area on the retina. $E_v$ is the energy of a 500 nm photon and can be calculated by $hc/\lambda$, where $h$ is the Planck constant, $c$ is the speed of light in vacuum, $\lambda =$ 500 nm is the photon's wavelength. Note that if the stimulus is broadband rather than just at 500 nm, additional calibration accounting for rhodopsin spectral sensitivity is required (5).

**Extraction of phase responses from the outer retina.** In phase-sensitive ORG measurements, we calculated the temporal phase change of a target layer with respect to a reference layer. For each pixel in the target layer, we selected a reference region from the reference layer centered at the same A-line. Each reference region comprised 5 adjacent A-lines (∼7.1 µm laterally). Before conducting spatial averaging across the reference region, we canceled out the systematic phase drift by self-referencing and removed arbitrary phase offset of the individual phase trace by referring to its pre-stimulus frames. Specifically, the systematic phase drift was canceled out by calculating the multiplication of the complex-valued OCT signal of the pixel of interest in the target layer and the complex conjugate of the signals in the selected reference region:

$$\tilde{I}_{tar/ref}(s,i) = \tilde{I}_{tar}(i)\tilde{I}^*_{ref}(s,i), \tag{S5}$$

where $\tilde{I}_{tar}(i)$ is the complex-valued OCT signal of the pixel of interest in the target layer, $i$ is the index of the frame number. $\tilde{I}_{ref}(s,i)$ represents signals in the corresponding reference region, and $s$ denotes the pixel index in the reference region. $\tilde{I}_{tar/ref}(s,i)$ is the pairwise self-referenced signal, where the pixel of interest in the target layer was ergodically referred to all pixels in its reference region. * represents complex conjugate.

To cancel out arbitrary phase offsets, each complex-valued signal trace was referred to its pre-stimulus frames,

$$\Delta\tilde{I}_{tar/ref}(s,i) = \tilde{I}_{tar/ref}(s,i) \cdot \frac{1}{N}\sum_{i=1}^{N}\tilde{I}^*_{tar/ref}(s,i), \tag{S6}$$

where $\Delta\tilde{I}_{tar/ref}(s,i)$ denotes time referenced signals, $N$ represents the number of frames acquired before the light stimulus. The complex-valued signals were subsequently averaged across pixels in the reference region, and the phase information was extracted from the averaged complex-valued signal:

$$\Delta\tilde{I}_{tar/ref}(i) = \frac{1}{S}\sum_{s=1}^{S}\Delta\tilde{I}_{tar/ref}(s,i), \tag{S7}$$

$$\Delta\phi(i) = \angle\Delta\tilde{I}_{tar/ref}(i), \tag{S8}$$

where $\Delta\phi(i)$ is one phase trace extracted from the pixel of interest in the target layer. $S$ is the total pixel number in the reference region, $\angle$ represents the calculation of argument.

We applied the same process to every pixel in the target layer to extract spatial-resolved temporal phase traces. Note that when averaging phase traces across different pixels, calculations were performed in the complex plane to avoid biases introduced by phase wrapping (6).

Phase traces can be converted into the OPL change ΔOPL by:

$$\Delta\text{OPL} = \frac{\lambda_0}{4\pi}\Delta\phi, \tag{S9}$$

or

$$\Delta\text{OPL} = -\frac{\lambda_0}{4\pi}\Delta\phi, \tag{S10}$$

where $\lambda_0$ is the central wavelength of the OCT system. When the reference layer was anterior to the target layer, the OPL change was calculated using Eq. (S9). Otherwise, it was calculated using Eq. (S10). This ensured that an increase in OPL change (ΔOPL) consistently represented an expansion between the target layer and the reference layer.

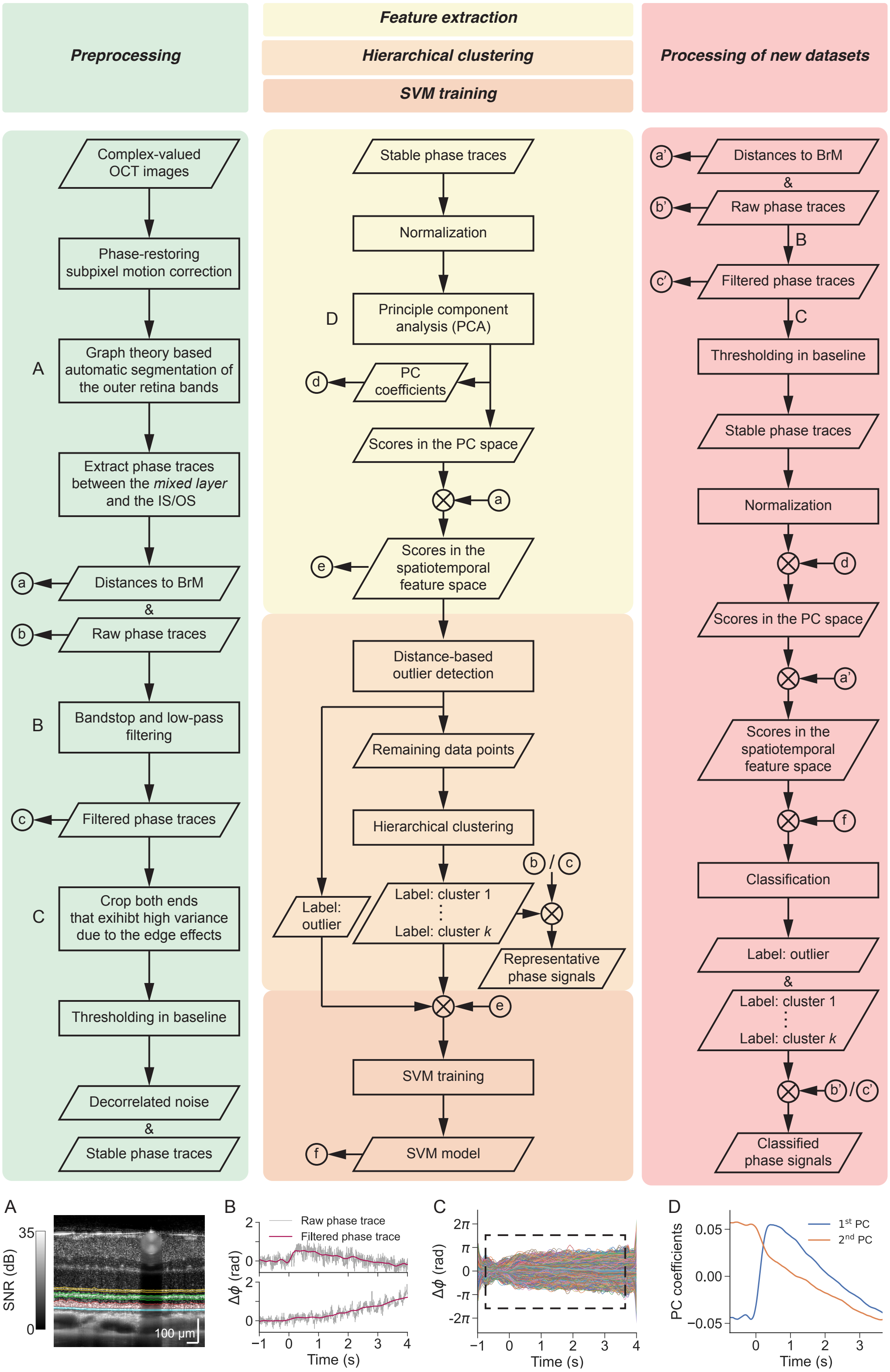


**Fig. S1.** Flow chart for classifying phase traces in the mixed layer. Phase traces extracted from the mixed layer were preprocessed and projected onto the spatiotemporal feature space. Distinct phase responses were identified using hierarchical clustering. A SVM was subsequently trained in the same feature space to facilitate the classification of new phase traces with the same criterion.

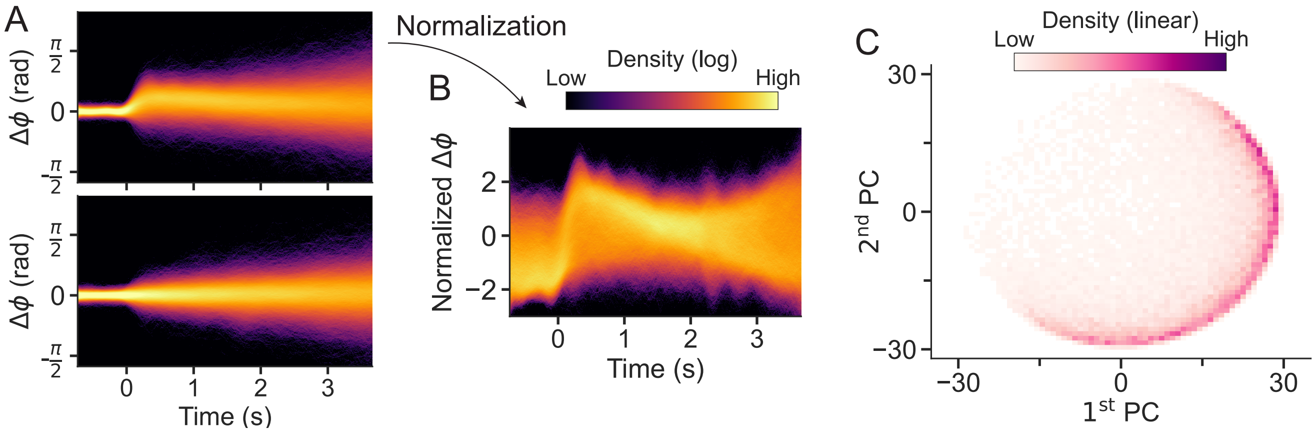


**Fig. S2.** Processing of phase traces and extraction of temporal signatures using principal component analysis. (A) The distribution density of phase traces that exhibited low variance prior to the light stimulus (before 0 s). When a flash was delivered to the retina (t = 0 s), function-associated phase responses were clearly observed (top panel), whereas without a light stimulus, phase traces were gradually decorrelated without any noticeable response pattern (bottom panel). (B) Each phase trace in the top panel of Fig. S2A was then normalized by subtracting its mean value and then divided by its standard deviation (SD). (C) The distribution of the normalized phase responses in the principal component (PC) space.

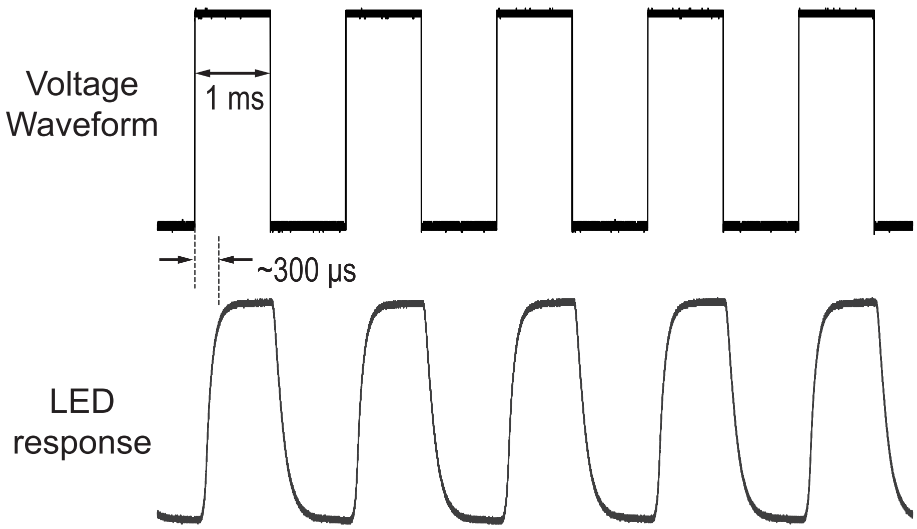


**Fig. S3.** LED's response waveform to trigger. Stimulus timing was delayed by approximately $300\,\mu s$, i.e., 6% of the B-scan frame acquisition time (5 ms).

**Table S1. Parameters used in the estimation of the hydraulic conductivity**

| Parameters | Value | Unit |
|---|---|---|
| Normal rodent plasma osmolarity (7), $\Pi_{\mathrm{rest}}/RT$ | 325 | mOsM |
| Maximum elongation*, $\Delta L$ | 33.6 | nm |
| Length of the rod OS, $L$ | 24.0 | $\mathrm{\mu m}$ |
| Surface area of the rod OS, $S_{\mathrm{ROS}}$ | 132 | $\mathrm{\mu m}^2$ |
| Cross-sectional area of the rod OS (1), $A_{\mathrm{ROS}}$ | 1.54 | $\mathrm{\mu m}^2$ |
| Initial water influx rate*, $J_w$ | 0.13 | $\mathrm{\mu m}^3{\cdot}\mathrm{s}^{-1}$ |

*: calculated with a refractive index of 1.41.

**Table S2. Animal usage**

| Test No. | Test Name | Number (Male) | Age |
|---|---|---|---|
| 1 | Dark adaptation | 12 (10) | 7-16 weeks |
| 2 | Light adaptation | 15 (10) | 6-12 weeks |
| 3 | Prolonged recording | 9 (6) | 6-12 weeks |
| 4 | Three-dimensional visualization | 7 (4) | 6-10 weeks |

**Table S3. Acquisition protocols**

| Test Number (refer to Table S2) | 1 | 2 | 3 | 4 |
|---|---|---|---|---|
| Acquisition Mode | Repeated B-scan | Repeated B-scan | Repeated B-scan | Repeated volumetric scan |
| Total Number of B-scans | 1000 | 1000 | 1375 | 1000 |
| Number of B-scans per Volume | N/A | N/A | N/A | 25 |
| Total time of acquisition (s) | 5 | 5 | 55 | 5 |
| Time interval between acquisitions (min) | 2 | N/A | 2 | 2 |
| Light/Dark adaptation | Dark adaptation | Light adaptation | Dark adaptation | Dark adaptation |
| Background intensity $(\mathrm{photons}/(\mathrm{\mu m}^2{\cdot}\mathrm{s}))$ | N/A | $6 \times 10^1$ - $6 \times 10^4$ | N/A | N/A |
| Flash duration (ms) | 1 | 1 | 1 | 1 |
| Flash bleach level (%) | 0.0019 - 0.28 | 0.28 | 0.18 & 0.26 | 0.10 |

**Movie S1.** ***En-face*** **temporal evolution of the outer segment (OS) and subretinal space (SRS) signals. Scale bar:** 200 μm.
